# Direct multiplex recombinase polymerase amplification for rapid detection of *Staphylococcus aureus* and *Pseudomonas aeruginosa* in food

**DOI:** 10.1101/2021.08.29.458066

**Authors:** Diem Hong Tran, Hau Thi Tran, Trang Nguyen Minh Pham, Huong Thi Thu Phung

## Abstract

Foodborne illness undermines human health by causing fever, stomachache and even lethality. Among foodborne bacterial pathogens, *Staphylococcus aureus* and *Pseudomonas aeruginosa* are of extraordinary significance which drive reasons of food and beverage poisoning in numerous cases. Today, PCR has been widely used to examine the presence of different foodborne pathogens. However, PCR requires specialized equipment and skillful personnel which limit its application in the field. Recently, there is an emerging of isothermal PCR methods in which the reactions occur at low and constant temperature, allowing their application in restricted-resource settings. In this work, multiplex Recombinase Polymerase Amplification (RPA) was used to simultaneously detect *S. aureus* and *P. aeruginosa* with high sensitivity and specificity. The limit detection of multiplex RPA was 10 and 30 fg/reaction of genomic DNAs of *S. aureus* and *P. aeruginosa*, respectively. Besides, the reaction time was reduced to only 25 minutes with a low incubation temperature of 39 °C. Markedly, multiplex RPA reactions succeeded to directly detect as low as 1 and 5 CFU/reaction of *S. aureus* and *P. aeruginosa* cells, respectively without the requirement of extracting DNA genome. Moreover, the multiplex RPA reliably detected the two foodborne bacteria in milk, fruit juice and bottled water samples. In general, the direct multiplex RPA described in this study is a rapid, simple, sensitive and efficient alternative tool that could be used to detect the presence of *S. aureus* and *P. aeruginosa* without the necessity of costly devices and high-trained staff.

## 1. Introduction

Most foodborne diseases are caused by bacterial pathogens, such as *Staphylococcus aureus, Pseudomonas aeruginosa, Salmonella enterica serovar Enteritidis*, and *Listeria monocytogenes* [1]. Accordingly, foodborne bacterial infections occur when the bacteria-contaminated food is consumed and the bacteria continue to develop in the stomach and intestines, generating a disease-causing infection. *S. aureus* is an opportunistic pathogen that can cause a range of life-threatening illnesses in animals and humans. Also, *S. aureus* is a major cause of food poisoning resulting from enterotoxin-contaminated food and drinks [2]. Meanwhile, *P. aeruginosa* is denoted by the World Health Organization as an indicator of drinking water quality [3]. *P. aeruginosa* can lead to long-term chronic diseases, particularly in patients admitted to the Intensive Care Unit [4]. Both *S. aureus* and *P. aeruginosa* can cause serious infections at extremely low concentrations [5]. Early detection with high sensitivity and specificity is therefore crucial to the treatment of *P. aeruginosa-* and *S. aureus*-derived diseases.

Traditional culture-based techniques for identification of bacterial pathogens typically require procedures of pre-enrichment, specific cultivation, and biochemical identification. Initial results thus need several days to verify a particular pathogenic microorganism, making those methods time-consuming and not practical for real-time application. Recently, newly pathogen-detecting molecular approaches have been intensively created and PCR-based methods are widely recognized as the gold-standard for diagnosing foodborne pathogens including *S. aureus* and *P. aeruginosa* [6-8]. These methods are able to detect a small number of bacterial cells. However, by virtue of requiring repeated thermal cycles with advanced equipment, complex processes and high costs, these techniques are limited in diagnosing food samples in the field with restricted resources. Therefore, new methods with elevated sensitivity and specificity that are quick, simple, easy to use, independent of complicated instruments have urgently been necessary.

Isothermal amplification is a current method for temperature-constant DNA amplification which provides an easy, rapid and economic approach to identify biological targets, particularly in less-equipped laboratories and at field application [9, 10]. Among these methods, a novel isothermal DNA amplification, namely recombinase polymerase amplification (RPA) has been introduced by scientists from ASM Scientific (Cambridge, UK) in 2006 [11] and rapidly improved for the molecular diagnosis of many infectious diseases including foodborne illness [12-15]. Later, ASM Scientific changed its name to TwistDx Ltd. and is still the only company that offers RPA kits on the market today. In 2017, Abbott acquired TwistDx and applied RPA to rapid diagnostic systems of infectious diseases. RPA uses a recombinase-primer complex, a strand-displacing polymerase, and single-stranded DNA binding proteins to replace the conventional thermal denaturation process of PCR, facilitating amplification of specific DNA products [16]. Compared with other isothermal amplification methods, RPA is the one whose operating principle and product are closest to PCR. Hence, its usage and applications also have many similarities and are easy to apply to laboratories that are familiar with PCR. Recently, combining with reverse-transcriptase (RT)-RPA, Xia *et al*. developed an one-pot, 30-min WEPEAR (whole-course encapsulated procedure for exponential amplification from RNA) protocol to diagnose SARS-CoV-2, showing a detection limit of 1 RNA copy/reaction [17]. Markedly, RT-RPA for SARS-CoV-2 detection was shown to possess the comparable sensitivity, specificity, and limit of detection (LOD) to those of RT-PCR [18]. These altogether indicate that the value of RPA technology is still being paid special attention nowadays.

Previously, RPA was successfully applied to rapidly detect *S. aureus* and *P. aeruginosa* [19, 20]. Nevertheless, different bacterial pathogens can usually co-infect food, thus quickly and accurately diagnose their presences at the same time will significantly save time, labor, and cost and bring more useful testing information as well. Consequently, in this study, we developed the multiplex RPA assay to simultaneously detect *S. aureus* and *P. aeruginosa* infected in drinking food. This assay is fast and simple and exhibits extremely high sensitivity and specificity. Especially, the assay is able to directly detect the bacterial cells without the requirement of DNA extraction in a contaminated water, fruit, and milk samples.

## 2. Material and method

### 2.1. Bacteria strains and reagents

A total of 7 common bacterial strains associated with dairy products and human health are listed in **Table 1**. The bacterial strains were cultured at 37 °C overnight. An RPA basic amplification kit including dried reagents, a rehydration buffer, and magnesium acetate was purchased from TwistDx (Cambridge, UK) for isothermal amplification of target sequences. Milk, fruit and bottled water were obtained from the local supermarket. Other chemical and necessary reagents were acquired from Sigma-Aldrich (USA).

**Table 1.**
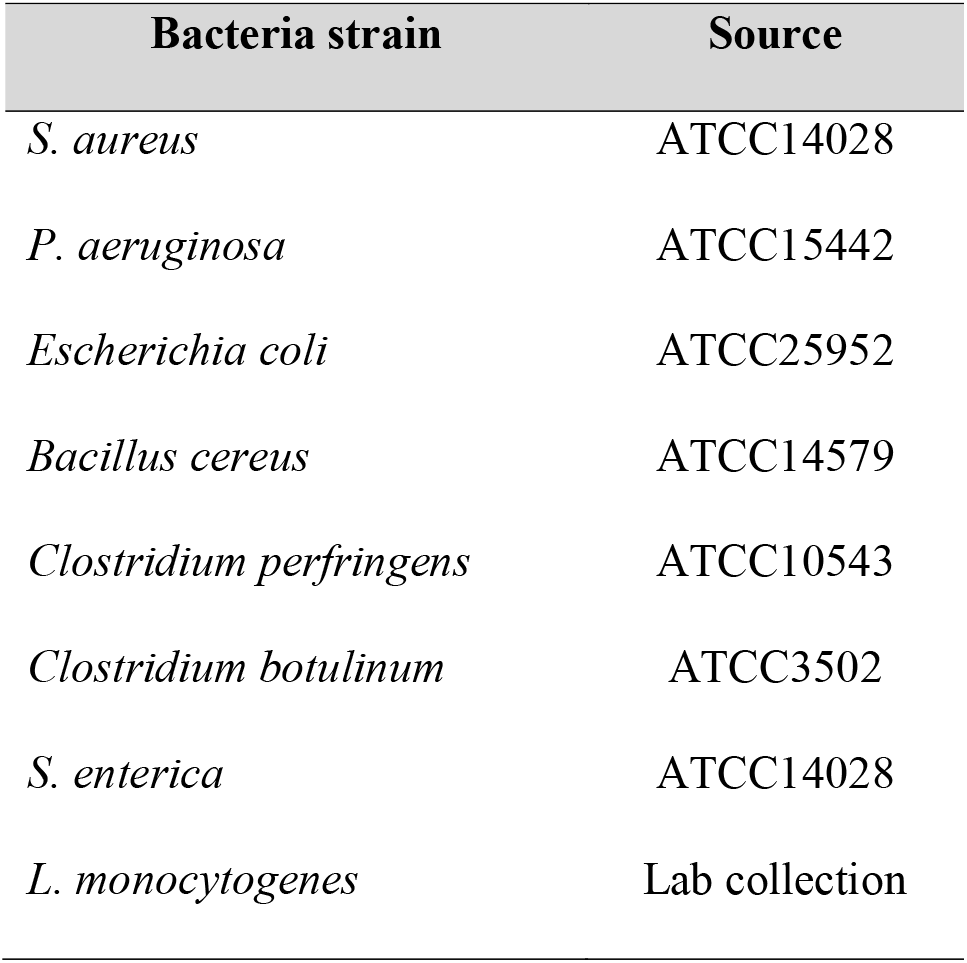
Bacteria strains used in this study.

### 2.2. DNA extraction and bacterial cells counting

The cells were incubated overnight in Luria Broth and the genomic DNAs were extracted using Cetyltrimethylammonium bromide (CTAB) method [21]. The concentration and quality of extracted genomic-DNAs were assessed by the optical density 260 nm and the ratio of 260/280 using a Genova Plus Spectrophotometer (Jenway, UK). DNA was stored at -20 °C until further use. The standard plate count method consisting of diluting a sample with sterile saline diluent until the bacteria are diluted enough to be counted accurately was used to calculate the number of cells. The final plates in the serial dilution should have between 30 and 300 colonies.

### 2.3. Primer design and screening

RPA primers were designed to specifically target the *S. aureus Sau-02* and *P. aeruginosa lasB* sequence regions, respectively. The Primer3 program was used to design the RPA primers as recommended by TwistAmp (TwistDX, UK). Totally, 2 primer sets were designed to target each of the *S. aureus Sau-02* and *P. aeruginosa lasB* sequence regions, respectively. Singleplex RPA assay was used to evaluate the amplification efficiency of designed primer sets. The primers were screened based on the brightness and specificity of the amplified product visualized by agarose gel electrophoresis. Finally, one primer set was selected for each target region (**Table 2**). All primers were synthesized by IDT (Singapore) and suspended in nuclease-free water to a concentration of 10 µM.

**Table 2.**
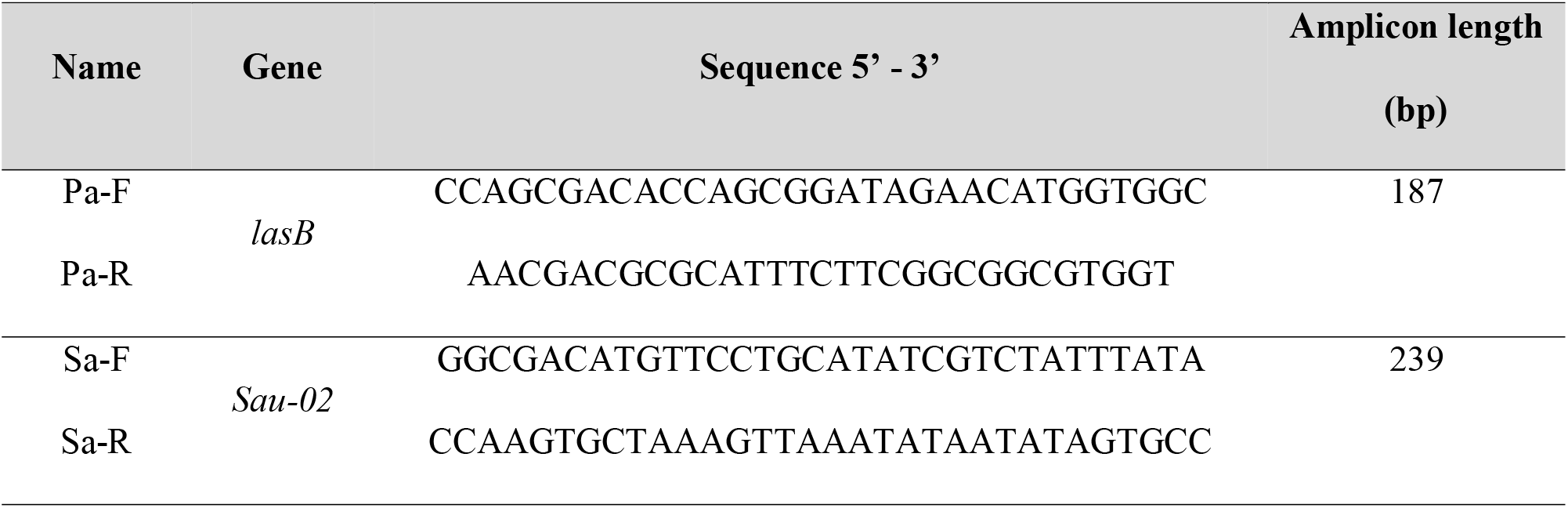
Primers used in RPA assay to detect *S. aureus* and *P. aeruginosa*.

### 2.4. Singleplex RPA assay

The RPA reaction contained 0.24 μM each of forward and reverse primer, 29.5 μL of rehydration buffer, 10.7 μL of nuclease-free water, 2.5 μL of template and a dried enzyme pellet. The reaction was initiated by adding 14 mM magnesium acetate. Next, the tubes were briefly centrifuged to mix all reagents and placed into the thermal incubator at the proper temperature for the indicated time. The RPA product was analyzed on a 2% agarose gel electrophoresis.

### 2.5. Multiplex RPA assay

Multiplex RPA reactions capable of amplifying two different targets were assembled by rehydrating an RPA enzyme pellet with 41.5-45.5 μl of a master mix depending on the volume of template. The master mix consisted of 29.5 μl of rehydration buffer, 7.2-11.2 μl of nuclease-free water, 0.24 μM each of *S. aureus* forward and reverse primers, and 0.24 μM each of *P. aeruginosa* forward and reverse primers. After rehydrating the enzyme pellet, the template and magnesium acetate (14 mM, final concentration) were added to initiate the reactions. For low copy-number template, the tubes were taken out after 4 minutes (min), vortexed, spun briefly and continued to be incubated in the heater until the time finished. Agarose gel electrophoresis was used to visualize the amplified products.

### 2.6. Optimization of Multiplex RPA Assay

Incubation time optimization of multiplex RPA assay was first conducted at different periods of time including 5, 15, 25, 35 and 45 min at 39 °C. The reactions were also incubated under isothermal conditions at the range of 35 to 42 °C for 30 min to define the optimal incubation temperature. Afterwards, agarose gel electrophoresis was used to analyze the amplification products.

### 2.7. Evaluation of specificity and limit of detection of multiplex RPA

The extracted genomic DNAs of several other bacteria relating to food poisoning were utilized as the template to evaluate the cross-reactivity of the RPA reactions. The *in-silico* PCR analysis using Genius Primer software (https://www.geneious.com/prime/) was also conducted for specificity evaluation of the multiplex RPA assay.

The DNA levels of *S. aureus* and *P. aeruginosa* were brought to the same concentration and a ten-fold serial dilution of genomic DNA was prepared to achieve the DNA concentration ranging from 5 to 10^4^ fg per reaction. One µl of each DNA dilution of each strain was used as the template for multiplex RPA reactions to determine the LOD values.

### 2.8. Evaluation of Direct Multiplex RPA Assay

To define the LOD of direct multiplex RPA test, cell cultures of *S. aureus* and *P. aeruginosa* were harvested and resuspended in nuclease-free water in order to produce the samples having the same cell concentration of each strain. The two cell solutions were then mixed with a ratio of 1:1 (v/v) and serially diluted in nuclease-free water to achieve the indicated cell concentration.

To evaluate the effectiveness of multiplex RPA assay for the direct detection of *S. aureus* and *P. aeruginosa* in drinking food, the cultured cells of *S. aureus* and *P. aeruginosa* collected were spiked into fresh milk, fruit juice or bottled water purchased from the local supermarket to produce drinking samples containing the indicated cell concentrations. One µl of the spiked samples was used as the template for the direct multiplex RPA tests.

## 3. Result

### 3.1. Optimization of Multiplex RPA Assay

The primer sets selected were utilized in the multiplex RPA reactions to detect both target sequences of *S. aureus Sau-02* and *P. aeruginosa lasB* (**Table 2**). The results shown in **Figure 1** indicate that the two primer sets successfully amplified both target sequences of *S. aureus* and *P. aeruginosa* simultaneously (**Figure 1, lane 3**). On the other hand, no product was observed in negative control reaction (**Figure 1, lane 4**).

**Figure 1.**
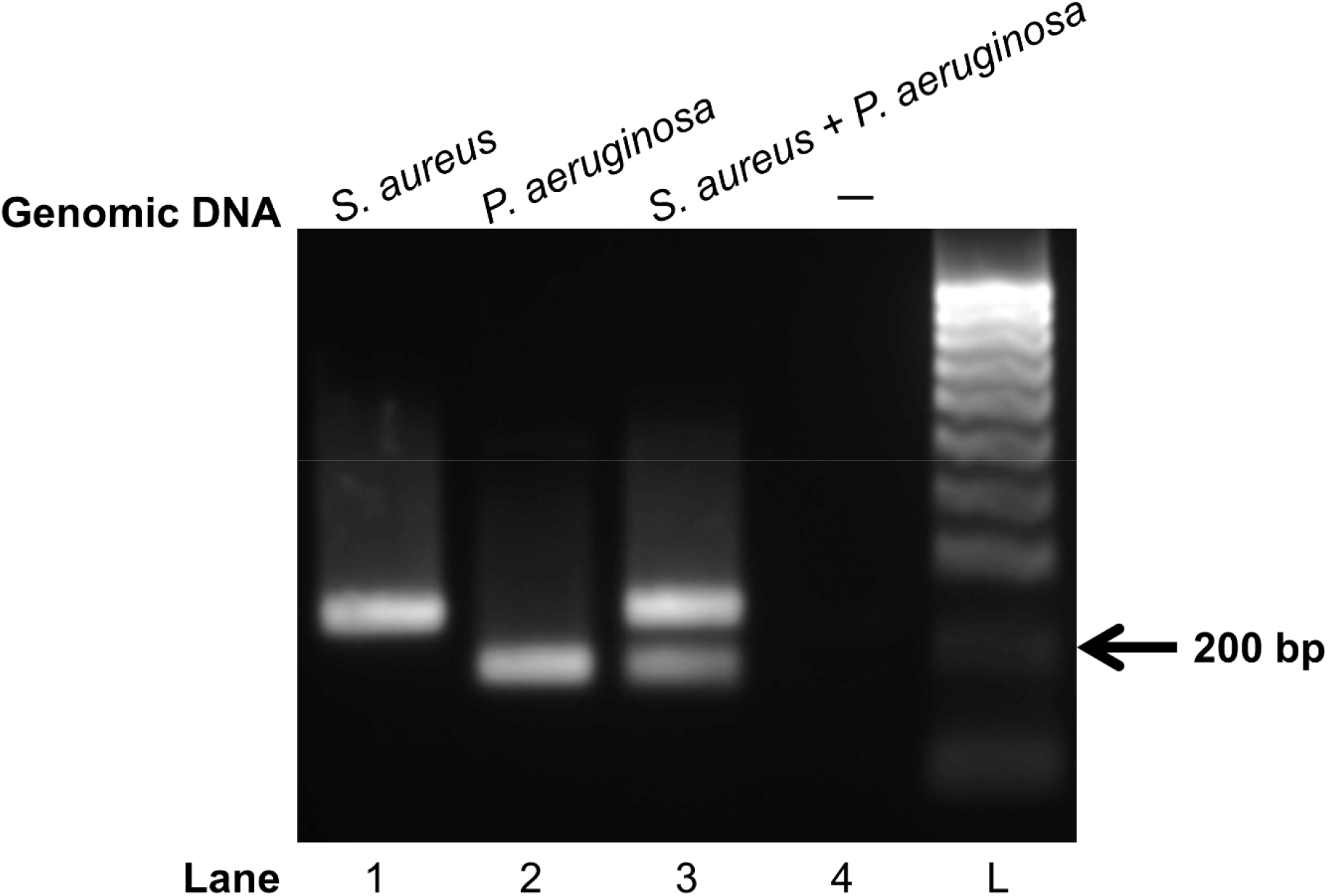
Multiplex RPA reactions to detect DNAs of *S. aureus* and *P. aeruginosa*. One ng of genomic DNAs of *S. aureus* and *P. aeruginosa* was used as the template for the singleplex and multiplex RPA reactions. The reactions were incubated at 39 °C for 30 min. Nuclease-free water was used as the negative control sample. The RPA products were analyzed on a 2% agarose gel electrophoresis. Abbreviation, L: DNA ladder.

Next, the multiplex RPA assay to simultaneously detect the DNA of *S. aureus* and *P. aeruginosa* was optimized regarding reaction time and incubation temperature. As shown in **Figure 2A lane 4**, after 15 min at 39 °C, the RPA amplified products could be observed by gel electrophoresis analysis. With longer incubation time, the band intensity of amplicon increased and saturated after 25 min (**Figure 2A, lane 3**). The reaction time was thus set for 25 min.

**Figure 2.**
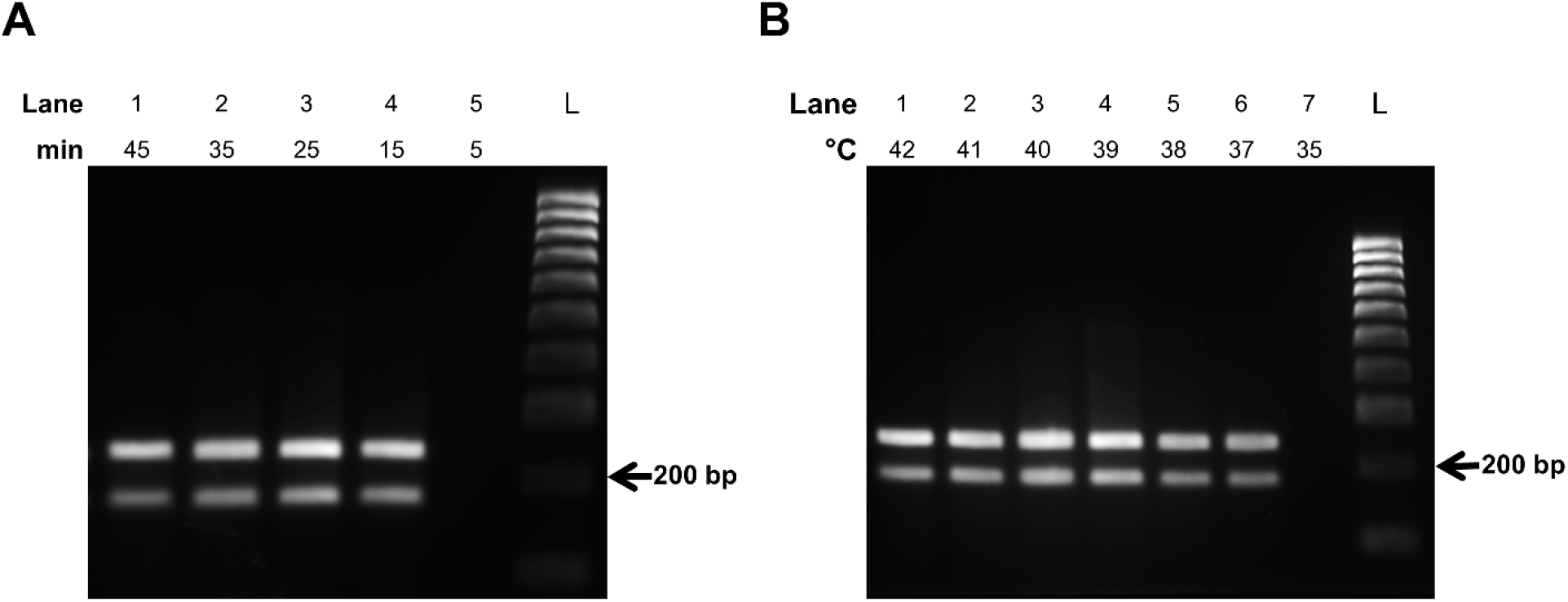
Optimization of the multiplex RPA reactions. The reactions contain one ng of each genomic DNA of *S. aureus* and *P. aeruginosa*. (A) The reactions were incubated at 39 °C for from 5 to 45 min. (B) The reactions were incubated at from 35 to 42 °C for 30 min. Abbreviation, L: DNA ladder.

Regarding incubation temperature, the optimal temperature of the multiplex RPA reaction was identified at 39 °C (**Figure 2B, lane 4**).

### 3.2. Specificity of Multiplex RPA Assay

To determine the specificity of multiplex RPA assay, extracted genomic DNAs of several other bacteria commonly associating with foodborne illness including *E. coli, L. monocytogenes, B. cereus, S. enterica, C. perfringens*, and *C. botulinum* were applied as templates for the multiplex RPA reactions. The results showed that only DNAs of *S. aureus* and *P. aeruginosa* strains generated amplified products while no amplicon was observed with other strains (**Table 3)**, indicating that there was no cross-reactivity among tested bacteria. The results of *in silico* PCR analysis shown in **Table 4** further demonstrated that the designed primer sets would not amplify any non-specific sequence of 20 different foodborne pathogens examined. Taken together, the obtained data support that the chosen primer sets possess a high specificity for *S. aureus* and *P. aeruginosa*.

**Table 3.** Cross-reactivity analysis of multiplex RPA assay detecting *S. aureus* and *P. aeruginosa*. One ng of genomic DNA of each strain was used for the multiplex RPA reaction. The reactions were incubated at 39 °C for 25 min.

| No. | Strain | RPA amplicon |
| --- | --- | --- |
| 1 | *S. aureus* | + |
| 2 | *P. aeruginosa* | + |
| 3 | *S. aureus* and *P. aeruginosa* | ++ |
| 4 | *E. coli* | — |
| 5 | *L. monocytogenes* | — |
| 6 | *B. cereus* | — |
| 7 | *S. enterica* | — |
| 8 | *C. perfringens* | — |
| 9 | *C. botulinum* | — |

**Table 4.** *In silico* PCR result (0-3 mismatch allowed in 3’-end)

| No. | Strain | *In silico* PCR product |  |
| --- | --- | --- | --- |
|  |  | *S. aureus*<br>primers | *P. aeruginosa*<br>primers |
| 1 | *S. aureus* | + | — |
| 2 | *P. aeruginosa* | — | + |
| 3 | *S. enterica* | — | — |
| 4 | *Salmonella typhi* | — | — |
| 5 | *C. perfringens* | — | — |
| 6 | *C. botulinum* | — | — |
| 7 | *Campylobacter fetus* | — | — |
| 8 | *Campylobacter jejuni* | — | — |
| 9 | *Campylobacter coli* | — | — |
| 10 | *Campylobacter upsaliensis* | — | — |
| 11 | *Campylobacter lari* | — | — |
| 12 | *Vibrio vulnificus* | — | — |
| 13 | *Vibrio cholera* | — | — |
| 14 | *Shigella dysenteriae* | — | — |
| 15 | *Shigella sonnei* | — | — |
| 16 | *Shigella flexneri* | — | — |
| 17 | *Cryptosporidium oocysts* | — | — |
| 18 | *Yersinia enterocolitica* | — | — |
| 19 | *Yersinia pestis* | — | — |
| 20 | *Yersinia pseudotuberculosis* | — | — |
| 21 | *B. cereus* | — | — |
| 22 | *E. coli* | — | — |

### 3.3. LOD of Multiplex RPA Assay

To identify the LOD value of multiplex RPA assay, a serial dilution of purified genomic DNAs of *S. aureus* and *P. aeruginosa* was used as templates for the RPA reactions. The results indicated that the corresponding product was not observed when the DNA template amount of *S. aureus* was lower than 10 fg/reaction (∼ 3.56 genome copies) (**Figure 3, lane 6**). Meanwhile, 30 fg/reaction (∼ 3.97 genome copies) was the lowest number of *P. aeruginosa* genomic DNA that was required for the relevant RPA amplicon formed successfully (**Figure 3, lane 5**). Thus, the LOD values of multiplex RPA assay for simultaneous detection of *S. aureus* and *P. aeruginosa* were determined at 10 and 30 fg/reaction of *S. aureus* and *P. aeruginosa* genomic DNAs, respectively.

**Figure 3.**
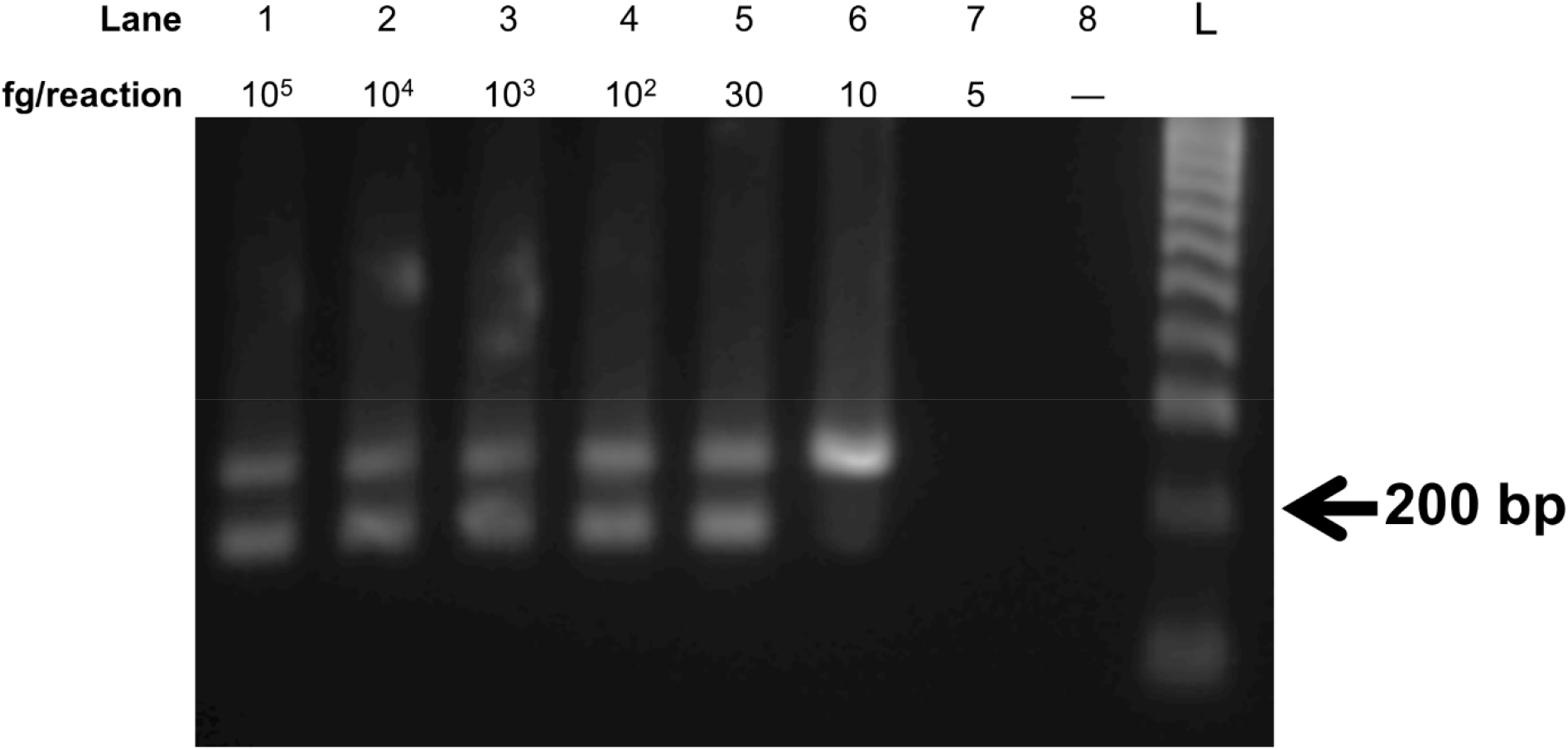
Evaluation of LOD of multiplex RPA assay simultaneously detecting *S. aureus* and *P. aeruginosa*. The LOD value of the multiplex RPA assay was evaluated using a serial dilution of *S. aureus* and *P. aeruginosa* genomic-DNA concentration ranging from 5 to 10^5^ fg/reaction. The reactions were incubated at 39 °C for 25 min. Abbreviation, L: DNA ladder.

### 3.4. Direct Multiplex RPA Assay

It was previously shown that RPA could directly detect different bacteria cells without the need of DNA extraction [22]. The amplification efficiency depends on the properties of the bacterial cell membrane and the initial membrane treatment methods [22]. In this study, *S. aureus* and *P. aeruginosa* cells were also used directly as the template for multiplex RPA reactions under the optimized condition for DNA-extracted template. The results showed that the multiplex RPA assay developed was capable of identifying the targeted DNA sequences when a mixture of *S. aureus* and *P. aeruginosa* cells was added to the reaction directly (**Figure 4**). Notably, the direct multiplex RPA assay was successful without a requirement of the cell-membrane treatment step. The LOD of direct multiplex RPA assay was thus evaluated by a serial dilution of the mixture containing *S. aureus* and *P. aeruginosa* cells. **Figure 4** indicates that the direct multiplex RPA can detect as low as 1 and 5 CFU/reaction of *S. aureus* and *P. aeruginosa* cells, respectively. There was no RPA amplicon generated at 0.5 CFU/reaction (**Figure 4, lane 6**). So, the LOD values of the direct multiplex RPA are determined at 1 CFU/reaction of *S. aureus* cells and 5 CFU/reaction of *P. aeruginosa* cells. This finding well agrees with the LOD values identified with genomic DNAs obtained above.

**Figure 4.**
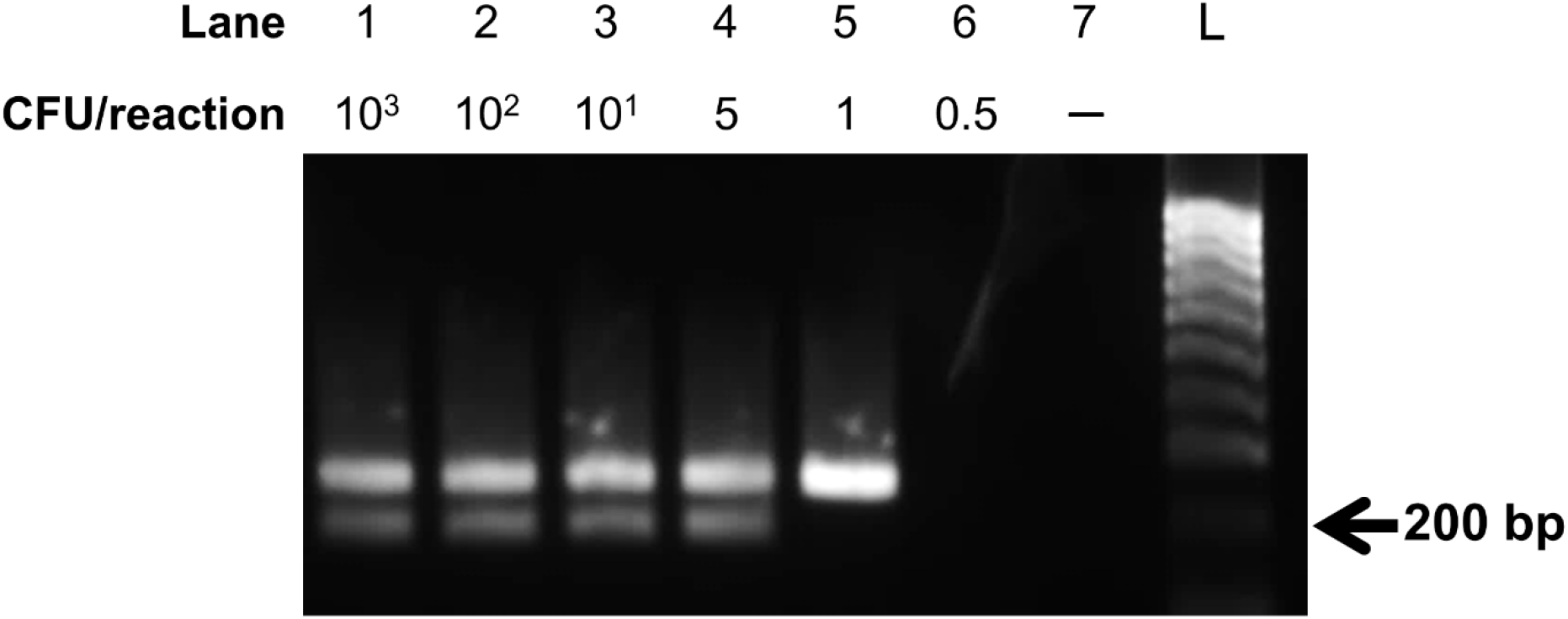
Evaluation of LOD of direct multiplex RPA detecting *S. aureus* and *P. aeruginosa* simultaneously. The cell mixture collected was diluted to concentrations ranging from 0.5 to 10^3^ CFU/reaction. One µl of each concentration was used for multiplex RPA reactions. The reactions were incubated at 39 °C for 25 min. Abbreviation, L: DNA ladder.

### 3.5. Direct detection of *S. aureus* and *P. aeruginosa* cells by RPA in food

The fresh milk, fruit juice and bottled water were used to evaluate the multiplex RPA assay for *S. aureus* and *P. aeruginosa* detection in food samples. To this end, the artificially contaminated drinking specimens were prepared by spiked with different concentrations of *S. aureus* and *P. aeruginosa* cells varying from 10^1^ to 10^3^ CFU/reaction. The simulated samples were then examined directly by the multiplex RPA reactions. Without the need for DNA extraction or cell-membrane treatment, the results revealed that *S. aureus* and *P. aeruginosa* cells could be successfully detected in food samples by the multiplex RPA assay developed (**Figure 5**).

**Figure 5.**
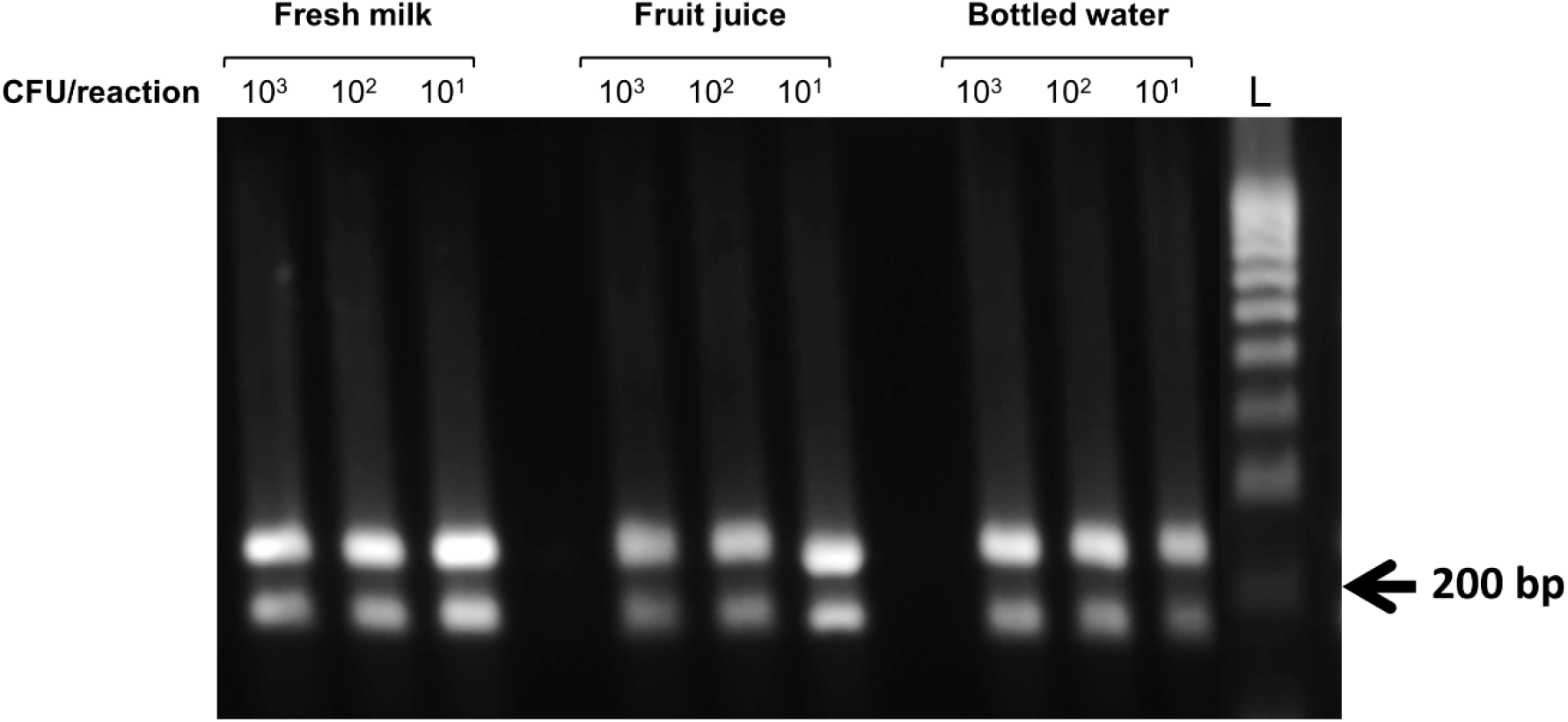
Performance of multiplex RPA assays for direct detection of *S. aureus* and *P. aeruginosa* cells in contaminated milk, fruit juice and bottled water. The RPA reactions were examined with the simulated food samples spiked with different concentrations of *S. aureus and P. aeruginosa* cells ranging from 10^1^ to 10^3^ CFU/reaction. One µl of each concentration was used for multiplex RPA reactions. The reactions were incubated at 39 °C for 25 min. Abbreviation, L: DNA ladder.

## 4. Discussion

The common pathogens *S. aureus* and *P. aeruginosa* were reported to cause various foodborne diseases such as typhoid fever, septicemia, and gastroenteritis, and even lethality [23]. Therefore, rapid, economic and reliable identification of *S. aureus* and *P. aeruginosa* has become particularly important. Numerous methods such as real-time PCR, enzyme-linked immunosorbent assays, and electrochemical biosensors have been developed to detect pathogenic bacteria in food [24-26]. However, because of the need for specialized equipment and time-consuming procedures, these methods are not ideal for implementation of broad diagnosis in the field. Recently, a variety of isothermal amplification techniques such as nucleic acid sequence-based amplification (NASBA), loop-mediated isothermal amplification (LAMP), helicase-dependent isothermal DNA amplification (HDA), RPA, cross-priming isothermal amplification (CPA), polymerase spiral reaction (PSR), etc have been established to replace PCR in detecting different infectious bacteria precisely and quickly on-site [27]. Nevertheless, multiple pathogens frequently coexist in food, hence the conventional detection of a single targeted DNA sequence cannot fulfill the need for multiple detection of different pathogens simultaneously. In contrast, a multiplex amplification reaction can simultaneously and rapidly detect multiple pathogens.

In this study, we developed the multiplex RPA assay for precise identification of the two foodborne pathogens simultaneously, namely *S. aureus* and *P. aeruginosa*. The multiplex RPA assay can be operated at a low constant temperature without a specialized instrument. It is well established that one of the most significant factors that enables the RPA assay to achieve the optimal efficiency and sensitivity is the reaction temperature. The recommended temperature for RPA reaction is between 35 and 42 °C [28]. Studies have shown that even at the environmental temperature including body temperature the RPA reaction can be conducted [29]. In our hand, the multiplex RPA amplicon could be produced at broad temperatures ranging from 37 to 42 °C. Accordingly, the optimal temperature for simultaneous detection of *S. aureus* and *P. aeruginosa* by the multiplex RPA assay established was determined at as low as 39 °C. Besides, the required incubation time was defined for just 25 min which is extremely shorter compared to a typical running time of PCR reaction. The low temperature and short incubation time of RPA reaction imply that the amplification efficiency is remarkable, pointing out that the RPA primers are well designed. In agreement, the designed primers also showed high selectivity for *S. aureus* and *P. aeruginosa*. The multiplex RPA assay can concomitantly detect 10 and 30 fg/reaction of genomic DNAs of *S. aureus* and *P. aeruginosa*, respectively. The success of multiplex RPA for simultaneous diagnosis of *S. aureus* and *P. aeruginosa* not only maintains the accuracy and sensitivity of the RPA technique but also reduces the number of reagents and operating steps required.

RPA was shown to possess a high impurity tolerance of the sample [30]. As expected, the direct addition of bacterial cells to the multiplex RPA reaction also induced the amplification process of the targeted sequences. It is worthy to mention that our direct multiplex RPA assay to identify *S. aureus* and *P. aeruginosa* cells does not require neither DNA extraction nor cell treatment prior to amplification reaction. The remarkable LOD determined at 1 and 5 CFU/reaction of *S. aureus* and *P. aeruginosa* cells, respectively further show that the direct multiplex RPA developed is extremely sensitive and robust. Importantly, the direct multiplex RPA assay was successfully applied on food samples without a requirement of sample processing, strengthening its high feasibility in real application at the field.

In general, this is the first study successfully established the direct multiplex RPA assay for simultaneous diagnosis of *S. aureus* and *P. aeruginosa*. The assay developed possesses the significant following advantages: (i) low incubation temperature required (39 °C) and short incubation time (less than 30 min), saving energy expense; (ii) multiple detection of two pathogens, decreasing the reagent amount and handling steps required; (iii) highly specific and sensitive; (iv) no need of DNA extraction and sample processing; and (v) easy to be applied for high-throughput diagnosis if needed. Consequently, further study should be performed to evaluate the clinical performance of the assay in realistic application.

## Acknowledgments

This work was funded by NTT Hi-Tech Institute, Nguyen Tat Thanh University under the grant number 2021.01.107

## Conflict of Interest

The authors declare no conflict of interest.

## Notes

### Competing Interest Statement

The authors have declared no competing interest.

